# Cerebral perfusion and metabolic response of astrocytes and neurons during locomotion

**DOI:** 10.1101/2024.12.23.630097

**Authors:** Alisa Tiaglik, Kseniia Morozova, Anna Fedotova, Alexey Brazhe, Li Li, Dan Liu, Ling-Qiang Zhu, Dmitry Bilan, Vladimir Oleinikov, Nadezda Brazhe, Alexey Semyanov

**Affiliations:** College of Medicine, Jiaxing University, Jiaxing, Zhejiang Province, 314001, China; Faculty of Biology, M.V. Lomonosov Moscow State University, Moscow, 119234, Russia; M.M. Shemyakin and Yu.A. Ovchinnikov Institute of Bioorganic Chemistry RAS, Moscow, 117997, Russia; Department of Medical Genetics, School of Basic Medicine, Tongji Medical College, Huazhong University of Science and Technology, Wuhan, Hubei 430030, China; Department of Pathophysiology, School of Basic Medicine, Tongji Medical College, Huazhong University of Science and Technology, Wuhan, Hubei 430030, China

## Abstract

The hemodynamic response links neural oxidative metabolism changes to increased cerebral blood flow during brain activity^1^. Although Roy and Sherrington introduced this concept over a century ago (1890), the exact cellular mechanisms remain unclear. This study demonstrates how local blood supply increases correlate with the metabolic response of individual brain cells during locomotion. Using Raman microspectroscopy, we observed an elevation in oxygen saturation levels (sO_2_) in cortical venules but not in arterioles, which were already near saturation. The increased sO_2_ in the venules was accompanied by vasodilation, indicating blood oversupply in the local brain area, a phenomenon known as functional hyperemia. We then analyzed the metabolic response of individual neurons and astrocytes to locomotion. In neurons, the levels of reduced cytochromes of *c* and *b* types [cyt *c,b* (Fe^2+^)] rapidly decreased at the onset of locomotion. This suggests an increase in the activity of the mitochondrial respiratory chain (electron transport chain, ETC) in response to heightened energy demands in these cells^2^. In contrast, because astrocytes rely less on oxidative phosphorylation for their energy metabolism than neurons^3^, we did not observe an initial decrease in reduced cytochromes in these cells. However, as locomotion continued, the cyt *c,b* (Fe^2+^) levels steadily increased in both cell types. In neurons, this led to a slow recovery from the initial drop, while in astrocytes, the increase exceeded baseline levels. Consequently, we observed an overload of electrons in the astrocytic ETC. The distinct responses of astrocytic and neuronal mitochondria to locomotion may reflect differences in the organization of the ETC in the two cell types^4^. Furthermore, astrocytes may shift to glycolysis under increased neuronal activity^5,6^. This difference also resulted in hydrogen peroxide (H_2_0_2_) production in astrocytic mitochondria but not neuronal mitochondria. Since H_2_0_2_ is a signaling molecule critical for cognitive function in the brain^7^, we speculate that astrocytic mitochondria act as signaling hubs during exercise.

## Introduction

Brain activity during physical exercise, such as locomotion, is linked to increased cerebral blood flow, which provides oxygen and energy substrates (glucose and fatty acids) necessary for ATP production^6^. This cerebral perfusion and oxygenation increase is a systemic response that includes elevated breathing and heart rates^8,9^. Additionally, the brain has specific mechanisms that regulate blood supply in active regions^6,10^. The region-specific increase in oxygenation, in response to heightened oxidative metabolism in neural cells, is known as the hemodynamic response. This concept is widely accepted for interpreting blood oxygen level-dependent (BOLD) functional magnetic resonance imaging (fMRI)^11,12^.

Astrocytes are crucial in regulating local blood supply^13,14^. During animal activity, these cells generate calcium transients^15,16^, which can trigger the synthesis or release of vasomotor effectors, such as metabolites of arachidonic acid, potassium, and nitric oxide (NO)^14,17,18^. Beyond regulating blood flow, astrocytes also provide metabolic support to neurons by shuttling lactate produced through astrocytic glycolysis, which is promoted by calcium elevations^5,6^.

The activation of glycolysis, alongside oxidative phosphorylation, defines the metabolic aspect of the hemodynamic response and can be associated with changes in the redox state of mitochondria in astrocytes and neurons.

In this study, we employed label-free Raman microspectroscopy (RM) for BOLD imaging (BOLD RM) in individual blood vessels while simultaneously imaging the mitochondrial redox state in astrocytes and neurons during locomotion in mice. The functional hyperemia induced by locomotion was associated with increased blood oxygenation in cortical venules but not in arterioles. This increased brain oxygenation was accompanied by opposite redox responses in neuronal and astrocytic mitochondria and astrocytic H₂O₂ production.

## Results

### RM in Mouse Brain during Locomotion

Label-free RM has been predominantly employed in geology, semiconductor, and materials sciences but has recently been applied to characterize biological samples *in vitro* and *in vivo*^19,20^. In this study, we used this technique for the first time to characterize metabolic processes in the mouse brain during locomotion.

Raman scattering differs from fluorescence because it does not rely on relatively long-living excited states, and it contrasts with Rayleigh (elastic) scattering as scattered light has a different wavelength than the incident light (Figures 1a)^19,21^. Raman scattering occurs on specific molecules based on their vibrational and rotational states, enabling the study of their concentrations, conformations, and redox states. We identified Raman spectral peaks specific to lipids, proteins, phenylalanine (Phe), reduced cytochromes types *c* and *b* [cyts *c,b* (Fe^2+^)], and oxy- and deoxyhemoglobin (oHb and dHb, respectively; Fig. 1a, b; Supplementary Tables 1 and 2, Supplementary Fig. 1). The 532 nm laser light was absorbed by *c*- and *b*-type hemes and scattered via resonance Raman mechanism, providing selective enhancement of the Raman scattering of cyts *c,b* (Fe^2+^), oHb, and dHb compared to other molecules in the brain tissue^22–24^.

**Figure 1:**
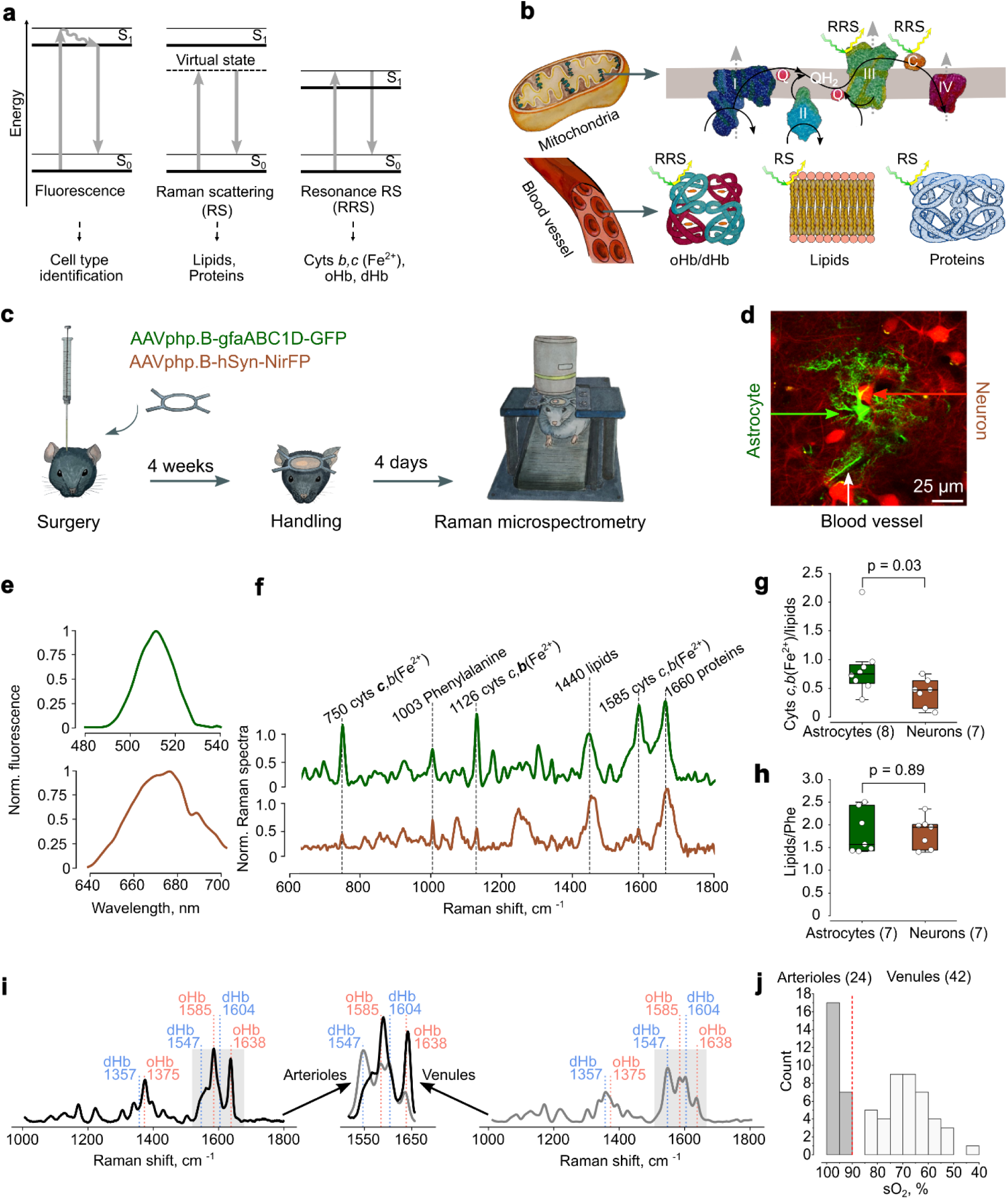
Raman and fluorescent imaging of astrocytes, neurons, and blood vessels in awake mice. **a**, The Jablonski diagram compares mechanisms of fluorescence, Raman scattering (RS), and resonance Raman scattering (RRS). S_0_ and S_1_ represent the ground and first excited singlet states, respectively. **b**, Fluorescence was utilized to identify cell types, RS was employed to quantify lipids and proteins, and RRS was used to quantify cyts *b,c* (Fe^2+^), oHb, and dHb. **c**, The experimental protocol involved stereotactic injection of two viral vectors into the somatosensory cortex (S1) for astrocytic expression of GFP (AAV.PHP.B-gfaABC1D-GFP) and neuronal expression of NirFP (AAV.PHP.B-hSyn-NirFP). Four weeks post-surgery, the animals were handled and adapted to the experimental setup for four days. Raman and fluorescent imaging were conducted on awake, head-fixed animals on a treadmill. **d**, Before RM, the expression of GFP in astrocytes and NirFP in neurons was assessed using two-photon imaging. Blood vessels were identified as relatively large structures devoid of fluorescent signals, surrounded by astrocytic endfeet. **e**, Fluorescence spectra obtained with a Raman microspectrometer corresponded to an astrocyte (*top*, green trace) and a neuron (*bottom*, brown trace). **f**, Raman spectra were obtained from an astrocyte (top, green trace) and a neuron (bottom, brown trace). The Raman peaks selected for further analysis are marked with the name of the molecule and the corresponding Raman shift value. **g**, Cyts *c,b* (Fe^2+^) peak at 750 cm^-1^ to lipid peak at 1440 cm^-1^ ratios in astrocytes and neurons are presented. **h**, Lipid peak to Phe peak ratios in astrocytes and neurons are presented. In **g** and **h**, circles represent individual data points; boxes indicate the median (Q1, Q3); whiskers represent 1.5 interquartile ranges (IQR). Statistical significance (p-values) was determined using the two-tailed Mann-Whitney test. The number of individual animals (N) is presented in brackets. One cell of each type was recorded per animal. **i**, Raman spectra for an arteriole (left) and a venule (right) are presented. The middle inset is an overlay of the greyed areas from the arteriole (black line) and venule (grey line) spectra. **j**, A histogram shows the distribution of blood vessels according to their sO_2_. The red dashed line indicates the boundary between arterioles (grey bars) and venules (white bars). The number of individual blood vessels (n) is presented in brackets, N = 7 animals.

We conducted multiplex fluorescence imaging and RM on the S1 somatosensory cortex of head-fixed mice running on a treadmill (Fig. 1c). Two adeno-associated virus (AAV) vectors were co-injected to express green fluorescent protein (GFP) in the cytosol of astrocytes and near-infrared fluorescent protein (NirFP) in the cytosol of neurons (Fig. 1c, d). The cells were identified by their respective fluorescence spectra obtained with 473 nm or 633 nm laser excitations before recording Raman spectra (Fig. 1e). The ratio of the Raman peak intensity of cyts *c,b* (Fe^2+^) to that of lipids was used to estimate the relative amount of cyts *c,b* (Fe^2+^) in mitochondria. We observed a significantly higher level of cyts *c,b* (Fe^2+^) in astrocytes than in neurons during animal quiescence (Fig. 1g). This finding suggests that ETC in astrocytes has a higher electron load than in neurons. Alternatively, the cyts *c,b* (Fe^2+^) ratio to lipids could also reflect differences in lipid content between the two cell types. However, the lack of statistically significant differences in lipids ratio to Phe ruled out this possibility (Fig. 1h).

The blood exhibited a distinctive Raman spectrum, characterized by peaks corresponding to the vibrations of heme bonds in oHb and dHb. This distinguished blood vessels and brain cells (Fig. 1i). The oHb ratio to dHb was utilized to calculate the oxygen saturation level (sO_2_). The distribution of blood vessels based on their sO_2_ revealed two pronounced peaks: arterioles (sO_2_: 96.8 (92.1, 100) %, n = 24) and venules (sO_2_: 68.9 (60.9, 73.3) %, n = 42, p < 0.001 for the difference with arterioles, two-sided Mann-Whitney test; Figure 1j) in the quiescent state of the animal.

### Effect of Locomotion on sO_2_ in Arterioles and Venules

Next, we investigated how locomotion (in this case, forcing an animal to run for 32 s) affects sO_2_ in individual blood vessels (Fig. 2a). Since the baseline sO_2_ levels were already close to saturation, we observed no significant increase in relative change, ΔsO_2_/sO_2_, in arterioles (Fig. 2b, c, e). In contrast, ΔsO_2_/sO_2_ rapidly increased in venules before slowly returning to initial values after locomotion (Fig. 2b, d, e). Notably, ΔsO_2_/sO_2_ was larger in venules with lower baseline sO_2_ (Fig. 2f). These findings suggest that the brain receives additional oxygen not because the incoming blood is more oxygenated but due to functional hyperemia.

**Figure 2:**
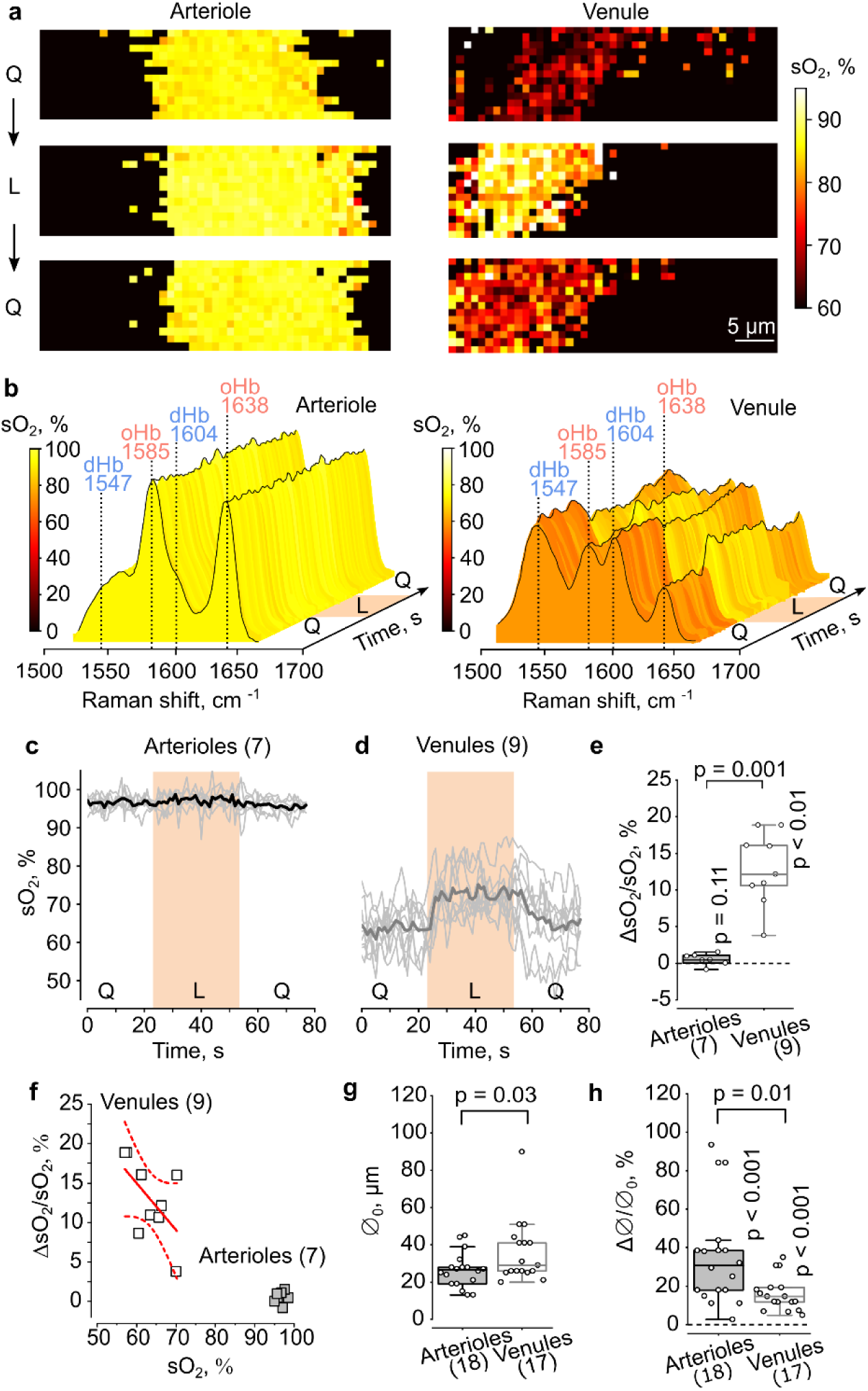
Blood oxygenation and functional hyperemia during locomotion. **a**, Raman mapping of sO_2_ in an arteriole (*left column*) and a venule (*right column*) during quiescence (Q), locomotion (L), and post-locomotion quiescence (Q) is shown. Color coding reflects the level of sO_2_. **b**, Raman spectra recorded in the arteriole (left) and venule (right) demonstrate changes in the peaks for oHb and dHb during quiescence (Q), locomotion (L), and post-locomotion quiescence (Q). The color coding corresponds to the level of sO_2_. **c**, Time courses of sO_2_ in arterioles during quiescence (Q), locomotion (L), and post-locomotion quiescence (Q) are presented. Thin grey lines represent individual arterioles; the thick black line indicates the mean trace. **d**, Similarly, time courses of sO_2_ in venules during quiescence (Q), locomotion (L), and post-locomotion quiescence (Q) periods are outlined. Thin grey lines denote individual venules; the thick grey line represents the mean trace. **e**, Summary data illustrate changes in sO_2_ during locomotion relative to pre-locomotion quiescence levels of sO_2_ (ΔsO_2_/sO_2_) in arterioles and venules. **f**, The dependence of ΔsO_2_/sO_2_ on pre-locomotion quiescence levels of sO_2_ in arterioles and venules is shown. The red line represents a linear fit for venules, while the dashed lines indicate the 95% confidence band. Pearson’s r = - 0.58 **g**, Summary data show the lumen diameter of arterioles and venules. **h**, Summary data show the change in lumen diameter during locomotion (Δ⌀) relative to pre-locomotion lumen diameter (⌀_0_) (Δ⌀/⌀_0_) in arterioles and venules. In **e**, **g**, and **h**, circles denote individual data points; boxes indicate the median (Q1, Q3); whiskers represent 1.5 IQR. p-values were calculated using the two-tailed Mann-Whitney test for comparisons between arterioles and venules and the two-tailed one-sample Wilcoxon Signed Rank test for significance of change. The number of individual blood vessels (n) is presented in brackets, N = 7 animals.

Increased blood flow in active brain areas is associated with vasodilation. Therefore, we measured the lumen diameter of blood vessels (⌀o) after identifying them as arterioles and venules by sO_2_ with Raman spectroscopy. Consistent with thicker walls, the median ⌀o was smaller in arterioles than in venules (Fig. 2g). Due to the larger number of smooth muscle cells, arterioles dilated significantly more than venules (Fig. 2h). This confirms that increased cerebral oxygenation is driven by enhanced blood supply to active brain areas during locomotion. The increased sO_2_ in venules suggests cerebral blood flow surpasses the levels needed to support heightened oxidative metabolism^25,26^.

### Metabolic Response of Astrocytes and Neurons

High metabolic demand during intense brain activity initially causes an undersupply of oxygen and nutrients, sequentially compensated by an oversupply^6,11^. We hypothesized that the metabolic activity of brain cells should also follow a biphasic pattern. To test this hypothesis, we measured the cyts *c,b* (Fe^2+^) level, which correlates with the number of electrons in the ETC of mitochondria and the efficiency of oxidative phosphorylation. We measured sO_2_ in blood vessels and cyts *c,b* (Fe^2+^) in astrocytic and neuronal mitochondria during locomotion (Fig. 3a).

**Figure 3:**
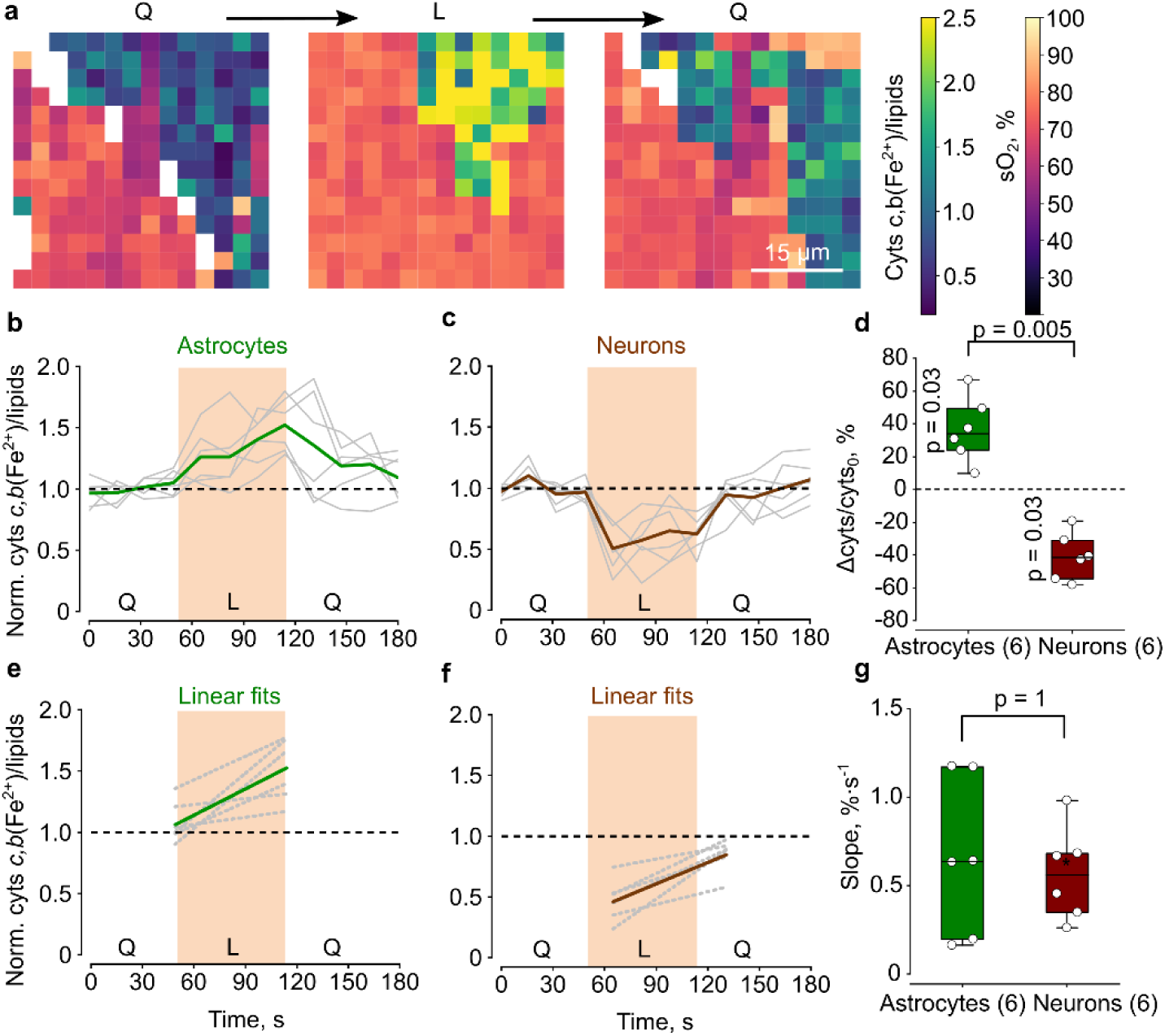
Opposite changes in the cyts *c,b* (Fe^2+^) levels in astrocytes and neurons during locomotion. **a**, Dual Raman mapping of sO_2_ in a blood vessel and cyts *c,b* (Fe^2+^) ratios to lipids in perivascular astrocyte was conducted during quiescence (Q), locomotion (L), and post-locomotion quiescence (Q). **b,c**, The time courses of cyts *c,b* (Fe^2+^) ratios to lipids in astrocytes (**b**) and neurons (**c**) normalized to cyts *c,b* (Fe^2+^) to lipids ratios during quiescence before locomotion. Thin grey lines indicate individual cells, while thick colored lines represent mean traces. **d**, Summary data showing changes in the cyts *c,b* (Fe^2+^) level during locomotion relative to the level in pre-locomotion quiescence (Δcyts/cyts_0_) is presented for astrocytes (green bar) and neurons (brown bar). **e**,**f**, Linear fits of changes in cyts *c,b* (Fe^2+^) ratios to lipids during locomotion in astrocytes (**e**) and neurons (**f**). Thin grey lines denote individual cells, and thick colored lines show mean traces. **g**, Summary data of the slopes of the fits presented in panels **e** and **f** for astrocytes (green bar) and neurons (brown bar). In panels **d** and **g**, circles denote individual data points; boxes indicate the median (Q1, Q3); whiskers represent 1.5 IQR. p-values were calculated using the two-tailed Mann-Whitney test for comparisons between astrocytes and neurons, and a two-tailed one-sample Wilcoxon Signed Rank test to assess the significance of changes in Δcyts/cyts_0_. The number of individual animals (N) is indicated in brackets. One neuron and one astrocyte were recorded per animal (n = N).

In astrocytes, the cyts *c,b* (Fe^2+^) level increased steadily during the locomotion period and slowly returned to baseline afterward (Fig. 3b). Conversely, in neurons, the cyts *c,b* (Fe^2+^) level sharply decreased at the onset of locomotion and slowly recovered to baseline during the remainder of the locomotion episode and after it ended (Fig. 3c). Thus, on average, the cyts *c,b* (Fe^2+^) level in astrocytic mitochondria increased during locomotion, while it decreased in neuronal mitochondria (Fig. 3d).

The steady increase of cyts *c,b* (Fe^2+^) in astrocytes and the gradual recovery in neurons may occur due to the increased blood supply to active brain regions. The additional blood enhances local brain oxygenation and delivers extra energy substrates, which supply electrons to the ETC. Notably, the rates of cyts *c,b* (Fe^2+^) growth in astrocytes and recovery in neurons were not significantly different, suggesting an equal supply of electrons to their ETC during the hemodynamic response (Fig. 3e, f).

### H_2_0_2_ Production by Astrocytes

The accumulation of electrons in the ETC, alongside increased tissue oxygenation, positions astrocytic mitochondria as a potential source of reactive oxygen species^27^. This phenomenon has been previously described in pathological conditions, such as during tissue reoxygenation after ischemia^28,29^. In this study, we examined whether such a mechanism is engaged under physiological conditions during locomotion. We employed mitochondria-tagged genetically encoded sensor HyPer7 to image reactive oxygen species in mitochondria of either astrocytes (Fig. 4a) or neurons (Fig. 4b). Indeed, we found a significant increase in reactive oxygen species in astrocytes but not in neurons (Fig. 4c).

**Figure 4:**
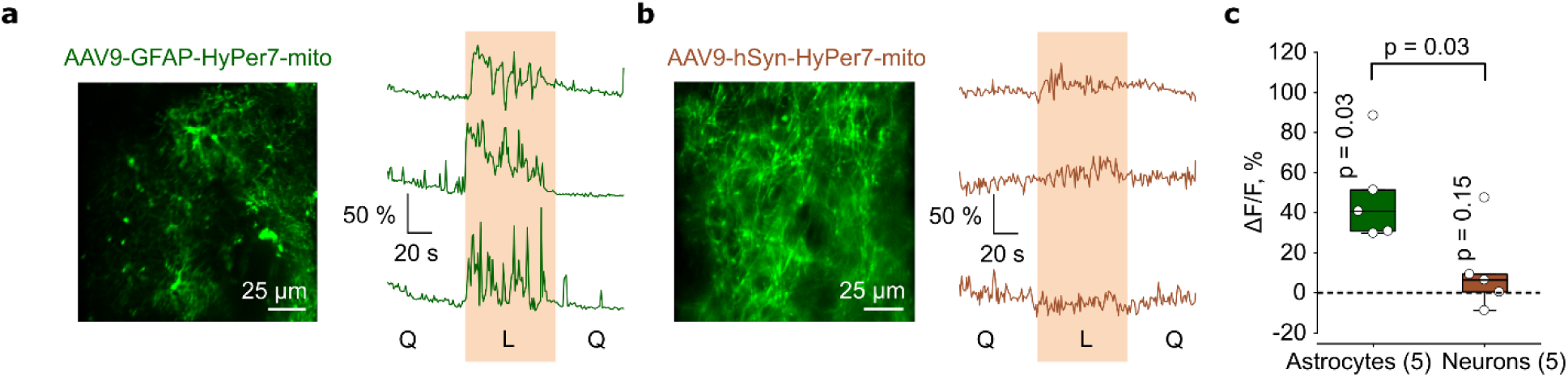
Locomotion is accompanied by H_2_O_2_ generation in astrocytes but not in neurons. **a**, On the left, a two-photon image shows astrocytes expressing the H_2_O_2_ sensor HyPer7 in their mitochondria following the injection of the AAV9-GFAP-HyPer7-mito viral vector. On the right, representative time courses of HyPer7 fluorescence in the mitochondria of three astrocytes during quiescence (Q), locomotion (L), and post-locomotion quiescence (Q) were measured using Raman microspectroscopy. **b**, On the left, a two-photon image of neurons expressing HyPer7 in their mitochondria following the injection of the AAV9-hSyn-HyPer7-mito viral vector. On the right, representative time courses of HyPer7 fluorescence in the mitochondria of three neurons during quiescence (Q), locomotion (L), and post-locomotion quiescence (Q) are shown. **c**, Summary data illustrate the changes in HyPer7 fluorescence during locomotion compared to its levels during pre-locomotion quiescence (ΔF/F) for astrocytes (green bar) and neurons (brown bar). Circles denote individual data points; boxes indicate the median (Q1, Q3); whiskers represent 1.5 IQR. p-values were calculated using the two-tailed Mann-Whitney test for comparisons between astrocytes and neurons and a one-tailed, one-sample Wilcoxon Signed Rank test for assessing the significance of changes in ΔF/F. The number of individual animals (N) is noted in brackets. Astrocytes and neurons were imaged in different animals, with one cell per animal (n = N).

## Discussion

We were the first to successfully perform multiplex imaging that combines fluorescence imaging and RM at a cellular resolution in awake animals. This approach allowed us to characterize the responses of three major components of the active brain milieu—blood vessels, astrocytes, and neurons—to locomotion^30^. Previously, Raman spectroscopy had been successfully applied to a variety of biological samples, including living cells in culture^23,31^, tissue slices^24,32,33^, and anesthetized animals^22,34,35^. However, changes in Raman spectra were often compared without quantifying specific molecules related to relevant cellular processes or the organism’s response. In our study, we employed RM to quantify the amounts of cyts *c,b* (Fe^2+^), lipids, Phe, oHb, and dHb during the awake state of the animals.

Our findings revealed that cortical astrocytes contain more than twice the amount of cyts *c,b* (Fe^2+^) compared to neurons. This result could indicate either a greater density of mitochondria (and total amounts of cytochromes) in astrocytes or a higher electron load in the ETC of these cells. A previous study demonstrated similar amounts of complex I in the mitochondria of both astrocytes and neurons, suggesting comparable amounts of cytochromes in these cells^36^. Therefore, our findings suggest an accumulation of electrons in the astrocytic ETC. This electron accumulation might be attributed to a different organization of the ETC in astrocytes compared to neurons^36,37^. Mitochondrial complexes are arranged in neurons into supercomplexes (respirasomes) to facilitate efficient electron transport and oxidative phosphorylation. In contrast, astrocytes maintain their ETC in a disassembled and energetically inefficient conformation, allowing electrons to linger longer at each complex, increasing the electron load and promoting reactive species’ generation. Indeed, higher hydrogen peroxide (H_2_O_2_) production in astrocytes compared to neurons has been reported in cell cultures^36^ and confirmed in our current *in vivo* experiments.

The varying amounts of cyts *c,b* (Fe^2+^) can be utilized for the label-free identification of astrocytes and neurons in brain tissue based on their Raman spectra. Additionally, RM can be employed for label-free measurements of oxygen saturation (sO_2_) in blood vessels, determined by the ratio of oHb to dHb (BOLD RM). This method enables the classification of blood vessels into arterioles and venules. Thus, RM can be beneficial when delivering fluorescent markers or viral transfections is not feasible, such as during human brain surgery or in optical brain-computer interfaces. Thus, our study can further advance the applications of Raman microspectroscopy in clinical research and as a diagnostic tool.

Brain activity during task performance or exercise increases the metabolic demand of brain tissue, which accelerates ATP production. This ATP fuels essential pumps, such as Na^+^/K^+^ ATPase or sarcoendoplasmic reticulum calcium ATPase, that restore transmembrane ionic gradients, the glutamate-glutamine cycle, and processes underlying plasticity in both neurons and glial cells^2,38^. A significant metabolic difference between neurons and astrocytes is their reliance on oxidative phosphorylation.

ATP production in mitochondria is vital for neuronal survival^3,6^. During locomotion, neuronal activity increases oxidative phosphorylation, which consumes available electron donors, which may explain the initial drop in neuronal cyts *c,b* (Fe^2+^) we observed. Enhanced blood supply delivers additional nutrients, gradually recovering cyts *c,b* (Fe^2+^) levels.

In contrast to neurons, astrocytes generate a significant amount of ATP through glycolysis^39,40^. Although astrocytes contain many mitochondria^41^, these organelles function as signaling hubs, producing reactive species necessary for sustaining cognitive performance^6,42^. Notably, we did not observe the initial drop in astrocytic cyts *c,b* (Fe^2+^) during locomotion; instead, there was a steady increase above baseline. This overload of the ETC with electrons was associated with the generation of H_2_O_2_, a versatile oxidant signaling agent^27^.

H_2_O_2_ interacts with iron-sulfur cluster proteins or the ionized form of cysteine residues (thiolate) on multiple redox-sensitive proteins involved in various physiological processes, including cell metabolism, phosphorylation cascades, transcriptional regulation, and cytoskeleton remodeling^27^. Astrocytic H_2_O_2_ regulates glucose utilization through the pentose-phosphate pathway, affecting glutathione metabolism and modulating the redox state of neurons^7^. A lack of astrocytic H_2_O_2_ production can lead to metabolic dysfunction and cognitive decline in mice^7^.

In conclusion, using multiplex fluorescent imaging, metabolic RM, and BOLD RM, we characterized the metabolic components of the hemodynamic response at the resolution of individual blood vessels, neurons, and astrocytes. The amount of neuronal cyts *c,b* (Fe^2+^) dropped, possibly reflecting the activation of oxidative phosphorylation. In contrast, the amount of astrocytic cyts *c,b* (Fe^2+^) gradually increased and correlated with H_2_O_2_ production in these cells. Since H_2_O_2_ has been suggested as a cognitive enhancer^7,36^, our results shed light on how exercise enhances brain function through the astrocytic component of the hemodynamic response. These findings also improve our understanding of BOLD fMRI recordings, which are not a direct measure of energy metabolism due to the mismatch between changes in blood flow and oxygen consumption^11^. BOLD RM also has greater sensitivity and selectivity for estimating blood oxygenation compared to widely used optical intrinsic signal imaging or near-infrared spectroscopy, whose signals can be ’contaminated’ by cyts *a,b,c* (Fe^2+^) components^43,44^. Finally, we demonstrate that RM can be employed as a label-free technique to identify individual cells and blood vessels in the brain and monitor their metabolic activity during behavioral tasks in experimental conditions or clinical studies (e.g., during surgeries or on biopsy materials)^24^.

## Methods

### Mice

All procedures followed the LASA ethical recommendations approved by the Institute of Bioorganic Chemistry ethical committee and the Laboratory Animal Ethics Committee of Jiaxing University. C57BL/6 mice (both sexes, 6-12 months old) were used for Raman confocal microspectroscopy and multiphoton imaging experiments. All animals were housed in groups of 2 to 3 in a 12-hour light-dark cycle and had access to food and water *ad libitum*.

### Surgery

Mice were anesthetized with a mixture of oxygen and isoflurane (1-1.5%) and placed in a stereotaxic frame (RWD, China). The skin over the head was removed, and a 3 mm craniotomy was performed above the somatosensory cortex (S1) using a dental drill (RWD, China). For co-labelling of astrocytes and neurons, 1 µl of AAV-PHP.B-hSyn-NirFP (5.1 x 10 ^12^ gc/ml^-1^, Eurogen Cat.# FP741) and AAV9-gfaABC1D-eGFP (2.42 x 10^11^ gc/ml^-1^, Addgene Cat.# 176861)^45^ were delivered to the coordinates (AP -2.3, ML +0.5, DV -0.8, -0.6, -0.4, -0.2). To detect H_2_O_2_ in the mitochondria of astrocytes or neurons, 1 µl of either AAV9-GFAP-HyPer7-mito (5.6 x 10^11^ gc/ml^-1^) or AAV9-hSyn-HyPer7-mito (1.2 x 10^12^ gc/ml^-1^) [Addgene 136470]^46^ was delivered to the same coordinates. AAV vectors were produced by the Viral Core Facility of IBCh RAS or VectorBuilder, Guangzhou, China.

Before and after each viral injection, the pipette tip was held for 5 minutes to ensure proper diffusion of viral particles in the region of interest. After the viral injection, a 5 mm cover glass (Thomas Scientific, USA) was sealed over the craniotomy site, and the head stage was affixed to the skull using a mixture of dental cement (Simplex Rapid Powder, Kemdent, UK) and cyanoacrylate glue (Cosmofen Ca-12, Germany). The experimental procedures started 4 weeks after surgery. The head stage was used to secure the mouse’s head under the objective of a Raman confocal microspectrometer (Renishaw, UK) or a multiphoton microscope (Femtonics, Hungary). The animals were habituated to the experimental setup and the environment before the experiments.

### Two-photon Imaging

A multiphoton fluorescence microscope (Femtonics FemtoSmart Dual, Hungary) with a 20x NA 1.0 objective was used to visualize astrocytes and neurons in the mouse somatosensory cortex (S1) in vivo. A Ti: Sapphire laser (Chameleon Ultra, USA) with a wavelength of 820 nm was used to excite the fluorescence of both GFP and NirFP. The emission was detected by GaAsP photomultipliers H10770PA-40 (Hamamatsu, Japan) after passing through a bandpass filter (520/60 nm for green light and 600/100 nm for red light). Fluorescence recordings were performed in galvo scanning mode with a size of the frame of 592 × 592 pixels and 0.5 µm per pixel.

### Treadmill

All experiments were conducted between 11 AM and 6 PM. Before the experiments, mice were habituated to the treadmill (https://doi.org/10.25378/janelia.24691311) and placed on the Raman microspectrometer motorized scanning table. The treadmill was cleaned between sessions using 70% ethanol and distilled water. Mice were head-fixed on the treadmill, and experimental sessions did not exceed 1-2 hours per mouse. All sessions were video recorded using an ELP 1.3-megapixel USB camera (Shenzhen Ailipu Technology Co., Ltd, China) at 20 frames per second. The video files were utilized for behavioral segmentation with the open-source program BORIS^47^. Raman spectra from blood vessels, astrocytes, and neurons were recorded during a quiescent state, stimulated locomotion, and post-locomotion quiescence. Stimulated locomotion was induced by running the treadmill belt at 15 cm/s.

### In Vivo Raman Microspectroscopy

Monitoring of local blood oxygenation and changes in the redox state of mitochondrial cytochromes in identified astrocytes and neurons of the somatosensory cortex (S1) of awake mice was conducted using a confocal Raman microspectrometer (InVia Qontor, Renishaw, UK) connected to a specially designed free-space upright microscope (Leica, Germany) with a 20x NA 0.4 objective. Raman spectra from blood vessels, astrocytes, and neurons were recorded with laser illumination (< 1 mW) at a wavelength of 532 nm, suitable for resonance Raman scattering at *c* and *b*-type hemes present in hemoglobin and cyts *c,b* (Fe^2+^).

### BOLD RM and Lumen Diameters of Blood Vessels

Raman spectra were recorded from individual vessels with an accumulation time of 1 s/spectrum during quiescence (60 s), stimulated locomotion (32 s), and post-locomotion quiescence (60 s). Raman peaks for dHb were located at 1357, 1547, and 1604 cm^-1^, while for oHb they were shifted to 1375, 1585, and 1638 cm^-1^, respectively (Supplementary Table 1, Supplementary Fig. 1)^22,48^. Linear baseline correction was performed based on the peak-free regions at 1500-1530 and 1680-1700 cm^-1^. After correction, the peaks at 1547 cm^-1^ (dHb) and 1585 cm^-1^ (oHb) were used to calculate sO_2_. Since the Raman cross-section of dHb is higher than that of oHb, a multiplier of 1.44 was used:

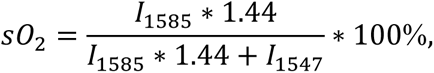

where I_1585_ and I_1547_ are the peak intensities of oHb and dHb, respectively. BOLD RM mapping was conducted at a spatial resolution of 1 µm/pixel using the “Map image acquisition” mode. We performed baseline correction on the Raman spectrum and calculated sO_2_ for each pixel to generate BOLD RM maps.

The lumen diameters of blood vessels were derived from sO_2_ profiles obtained from linescans across the vessels. The boundaries of the vessels were defined when sO_2_ dropped below 10% of the sO_2_ level in the center of the vessels.

### Measurement of Cyts *c,b* (Fe^2+^) in Astrocytes and Neurons

Astrocytes and neurons were identified using a Raman microspectrometer based on their fluorescence spectra of GFP and NirFP, respectively. The GFP fluorescence in astrocytic cytoplasm was excited using a 473 nm laser (grating 1200 lines/mm), while NirFP fluorescence in neuronal cytoplasm was excited with a 633 nm laser (grating 1200 lines/mm). After identifying the cell types, we recorded Raman spectra using a 532 nm laser (grating 1800 lines/mm) during quiescence (60 s), locomotion (60 s), and post-locomotion quiescence (60 s). The accumulation time for each Raman spectrum was 15 s. The reporter proteins GFP and NirFP did not interfere with Raman scattering excited by the 532 nm laser.

The Raman spectra were analyzed using custom-made software called Pyraman (https://github.com/abrazhe/pyraman). Baseline subtraction was performed for each spectrum, defined as a cubic spline interpolating a set of knots. The number and x-coordinates of these knots were selected manually, ensuring they were outside any informative peaks in the spectra. Once established, the number and x-coordinates of the knots remained constant for all spectra in the study. The y-coordinates of the knots were defined separately for each spectrum, representing 5-point neighborhood averages of spectrum intensities around the user-specified x-position of each knot. The parameters for baseline subtraction were selected after processing approximately 30 spectra from various astrocytes and neurons.

Following baseline correction, we obtained the intensities of the peaks at 750, 1003, and 1440 cm^−1^ (Supplementary Table 2): The 750 cm^−1^ peak corresponds to cyts *c,b* (Fe^2+^), with the majority contribution from *c*-type cytochromes, the 1003 cm^−1^ peak corresponds to Phe residues in proteins, the 1440 cm^−1^ peak corresponds to lipids^49^. We used the Raman peak intensities ratio of lipid content to Phe (I_1440_/I_1003_) and the cyts *c,b* (Fe^2+^) ratio to lipids (I_750_/I_1440_) in astrocytes and neurons. Cyts *c,b* (Fe^2+^) mapping was performed similarly to sO_2_ mapping but at a spatial resolution of 3 µm/pixel.

### H_2_O_2_ Measurement in Mitochondria of Astrocytes and Neurons

Monitoring H_2_O_2_ generation in astrocytes or neurons was carried out using the genetically encoded sensor HyPer7, expressed in the mitochondrial matrix of astrocytes or neurons within awake mice’s somatosensory cortex (S1). Cells were identified with the Raman microspectrometer by detecting HyPer7 fluorescence excited with a 473 nm laser at a power of 0.1 mW. Fluorescence spectra were collected using a 1200 lines/mm grating during quiescence, locomotion, and post-locomotion quiescence. After baseline correction, we estimated each spectrum’s maximum fluorescence intensity at 515 nm.

## Data availability

Datasets generated or analyzed during the current study are available on request.

## Code availability

Raman spectra were analyzed with the open-source software Pyraman, available at https://github.com/abrazhe/pyraman.

## Acknowledgments

Experiments conducted using Raman microspectroscopy received funding from RSF grant 23-44-00015 awarded to NB. The software development for analyzing Raman spectra, time series, and images was supported by RSF grant 23-44-00103 awarded to AB. AS’s work is supported by the Starlit South Lake Leading Elite Program 2022A 100001. The research conducted by L.Q.Z and D.L. was funded by the National Natural Science Foundation of China, with grant 82325017 awarded to L.Q.Z and grant 82261138555 awarded to D.L. The authors are grateful to Dr. Grigory Martynov for his help with building the treadmill.

## Authors’ contributions

NB, AS, LL, DL, and LQZ conceptualized the study and designed the experiments. AT, KM, and AF collected the data with input from NB, AS, VO, and DB. DB supplied a viral vector for cell-specific expression of HyPer7. AT, KM, AF, NB, and AS analyzed and interpreted the data. AB developed methods and software for analyzing Raman spectra, time series, and maps. AS, NB, and AT wrote the manuscript with input from all other authors.

---

**Supplementary Figure 1.**
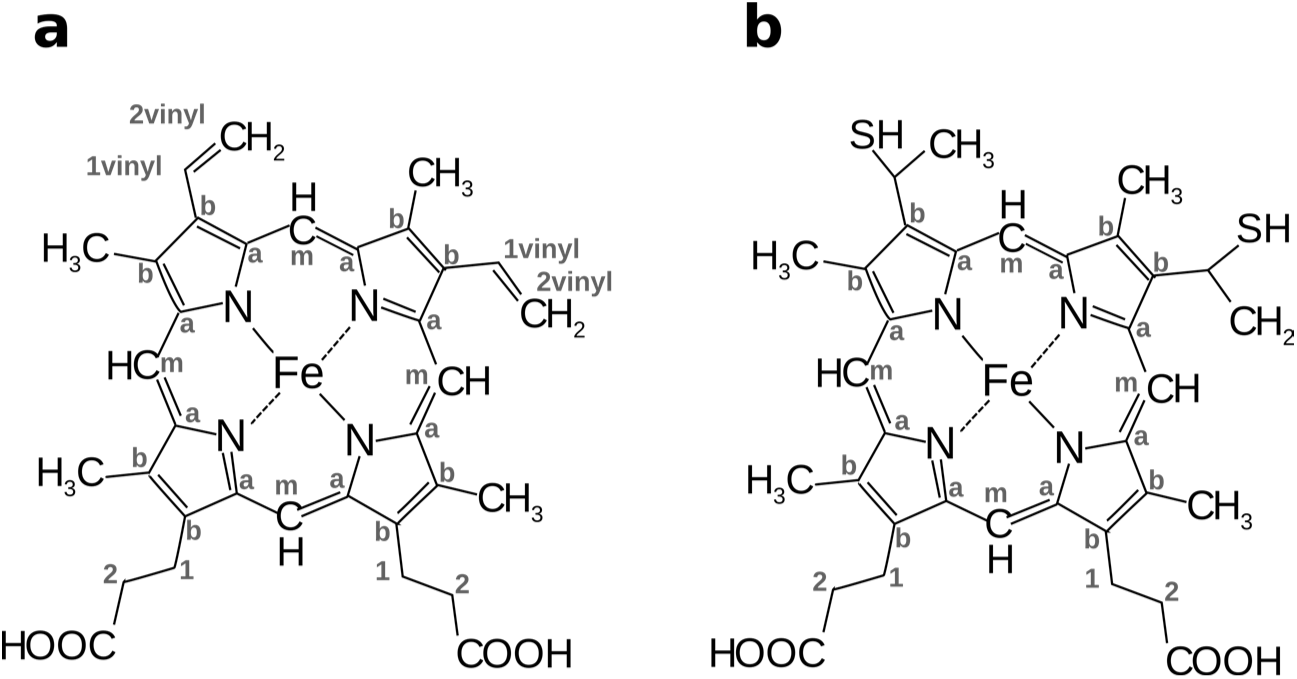
Structural formulas of *b*-type heme (a) and *c*-type heme (b) with the numeration of *C* atoms.

**Supplementary Table 1.**
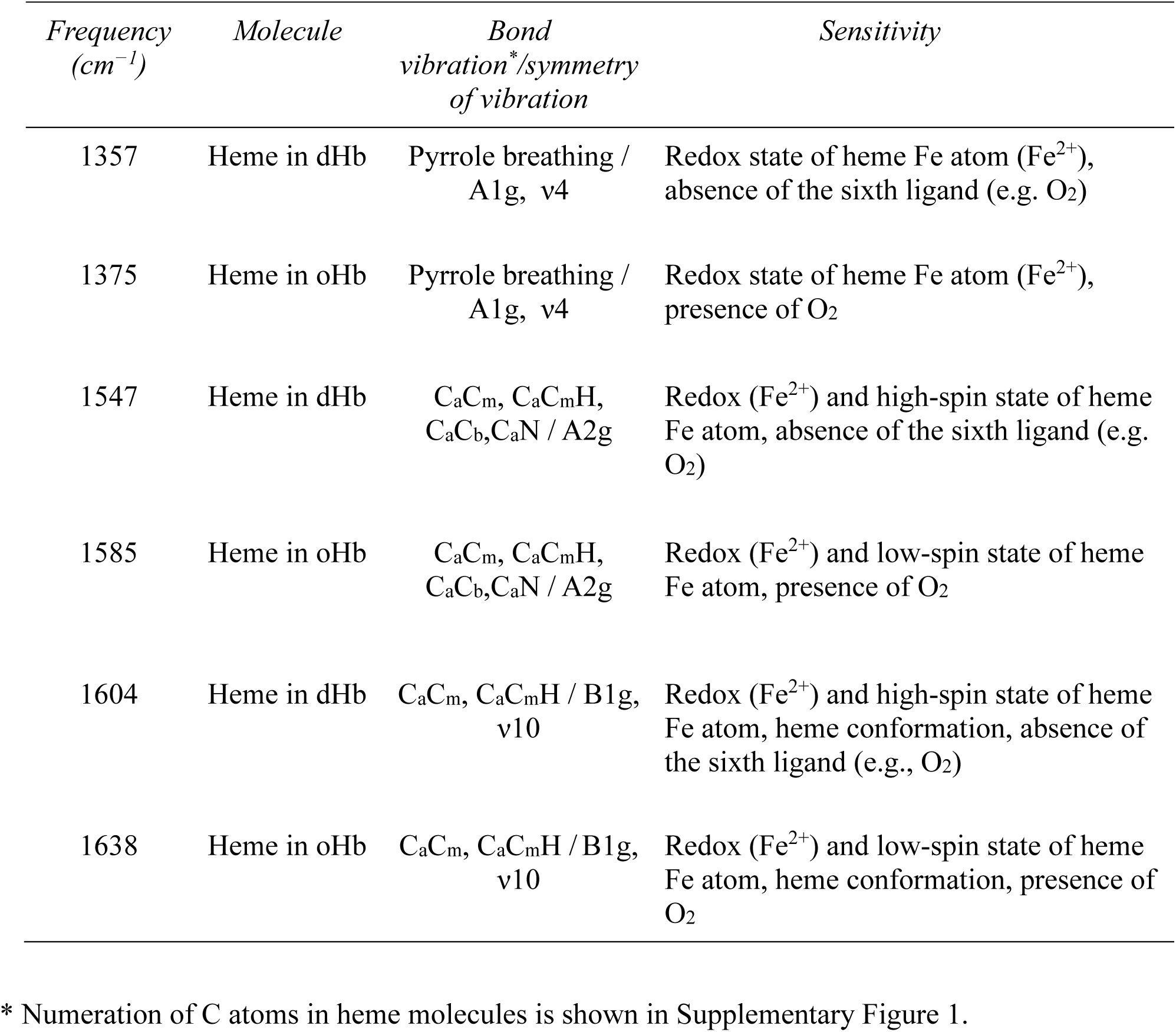
Assignment of peaks in Raman spectra of blood vessels in mouse cortex *in vivo (based on ^1-3^)*.

**Supplementary Table 2.** Assignment of peaks in Raman spectra of astrocytes and neurons in mouse cortex in vivo (based on ^4-9^).

| *Frequency*<br>*(cm<sup>-1</sup>)* | *Molecule* | *Bond*<br>*vibration\*/symmetry of vibration* | *Sensitivity* |
| --- | --- | --- | --- |
| 750 | Heme in reduced cytochromes *c*, *b* (Fe<sup>2+</sup>) | Heme breathing / B1g, v15 | Redox state of heme; the peak intensity is 10<sup>2</sup>-10<sup>3</sup> times higher for reduced cytochromes than for oxidized forms; the peak intensity is higher for reduced *c*-type cytochromes than for *b*-type |
| 1003 | Phenylalanine | Phe residue | Relative amount of protein |
| 1126 | Heme in reduced cytochromes *c*, *b* (Fe<sup>2+</sup>) | Heme methyl radical -CH<sub>3</sub> / B1g, v5 | Redox state of heme; the peak intensity is 10<sup>2</sup>-10<sup>3</sup> times higher for reduced cytochromes than for oxidized forms; the peak intensity is higher for reduced *b*-type cytochromes than for *c*-type |
| 1440 | Lipids | C-C and -CH<sub>2</sub> | Peak intensity depends on the relative amount of lipids |
| 1585 | Heme in reduced cytochromes *c*, *b* | CaC<sub>m</sub>, CaC<sub>m</sub>H, CaC<sub>b</sub>, CaN / A2g | Redox state of heme; the peak intensity is 10<sup>2</sup>-10<sup>3</sup> times higher for reduced cytochromes than for oxidized forms |
| 1660 | Proteins | C-N, peptide bond | Peak position depends on the secondary protein structure and the relative amount of protein. |

\* Numeration of C atoms in heme molecules is shown in Supplementary Figure 1.

